# Redefining Nephrotic Syndrome in Molecular Terms: Outcome-associated molecular clusters and patient stratification with noninvasive surrogate biomarkers

**DOI:** 10.1101/427880

**Authors:** Laura H. Mariani, Sean Eddy, Sebastian Martini, Felix Eichinger, Brad Godfrey, Viji Nair, Sharon G. Adler, Gerry B. Appel, Ambarish Athavale, Laura Barisoni, Elizabeth Brown, Dan C. Cattran, Katherine M. Dell, Vimal Derebail, Fernando C. Fervenza, Alessia Fornoni, Crystal A. Gadegbeku, Keisha L. Gibson, Deb Gipson, Lawrence A. Greenbaum, Sangeeta R. Hingorani, Michelle A. Hlandunewich, John Hogan, Larry B. Holzman, J. Ashley Jefferson, Frederick J. Kaskel, Jeffrey B. Kopp, Richard A. Lafayette, Kevin V. Lemley, John C. Lieske, Jen-Jar Lin, Kevin E. Myers, Patrick H. Nachman, Cindy C. Nast, Alicia M. Neu, Heather N. Reich, Kamal Sambandam, John R. Sedor, Christine B. Sethna, Tarak Srivastava, Howard Trachtman, Cheryl Tran, Chia-shi Wang, Matthias Kretzler

## Abstract

A tissue transcriptome driven classification of nephrotic syndrome patients identified a high risk group of patients with TNF activation and established a non-invasive marker panel for pathway activity assessment paving the way towards precision medicine trials in NS.

**Abstract:** Nephrotic syndrome from primary glomerular diseases can lead to chronic kidney disease (CKD) and/or end-stage renal disease (ESRD). Conventional diagnoses using a combination of clinical presentation and descriptive biopsy information do not accurately predict risk for progression in patients with nephrotic syndrome, which complicates disease management. To address this challenge, a transcriptome-driven approach was used to classify patients with minimal change disease and focal segmental glomerulosclerosis in the Nephrotic Syndrome Study Network (NEPTUNE). Transcriptome-based classification revealed a group of patients at risk for disease progression. High risk patients had a transcriptome profile consistent with TNF activation. Non-invasive urine biomarkers TIMP1 and CCL2 (MCP1), which are causally downstream of TNF, accurately predicted TNF activation in the NEPTUNE cohort setting the stage for patient stratification approaches and precision medicine in kidney disease.

## Introduction

Nephrotic syndrome (NS) refers to a glomerular disease with a shared clinical presentation, which is marked by proteinuria, hypoalbuminemia, hyperlipidemia and edema which can ultimately lead to kidney failure. Several underlying diseases can result in this constellation of symptoms, including the primary glomerular diseases of minimal change disease (MCD) and focal segmental glomerulosclerosis (FSGS), currently classified by the descriptive pattern of injury seen on kidney biopsy. Although these primary glomerular diseases are categorized as distinct histopathologic categories, they likely result from heterogeneous biological processes given the person to person variability in disease onset, rates of progression and response rates to various immunosuppressive therapies (*1*). Currently, diagnostic, prognostic and therapeutic decisions are based on these histopathologic categories and routine clinical parameters (e.g. serum creatinine and urine protein) that do not account for the heterogeneity of the biological antecedents. Because of the imprecise diagnosis within the current descriptive disease classification, molecularly targeted treatments for these diseases are not routinely available, and the interpretation of results from observational studies and clinical trials of therapeutic agents, which enroll a heterogeneous population of NS patients, are difficult to interpret. In such studies, it is often observed while the overall trial reads out negative, a small subset of patients respond well to the trialed therapy (*2–6*), yet pre-treatment predictors of response are not available.

Advances in biomedical research allow for capture of high-dimensional data across the genotype-phenotype continuum from patients under routine clinical care and can serve as a platform for implementation of precision medicine within NS (*7*). This approach utilizes large scale data integration across multiple data domains paired with deep clinical phenotype to establish a disease classification which is based in molecular causes as well as clinical presentation. Ultimately, the overarching goal of this approach is to assign targeted treatment based on these refined diagnostic categories which can reliably be identified using non-invasive markers (Figure 1).

**Figure 1:**
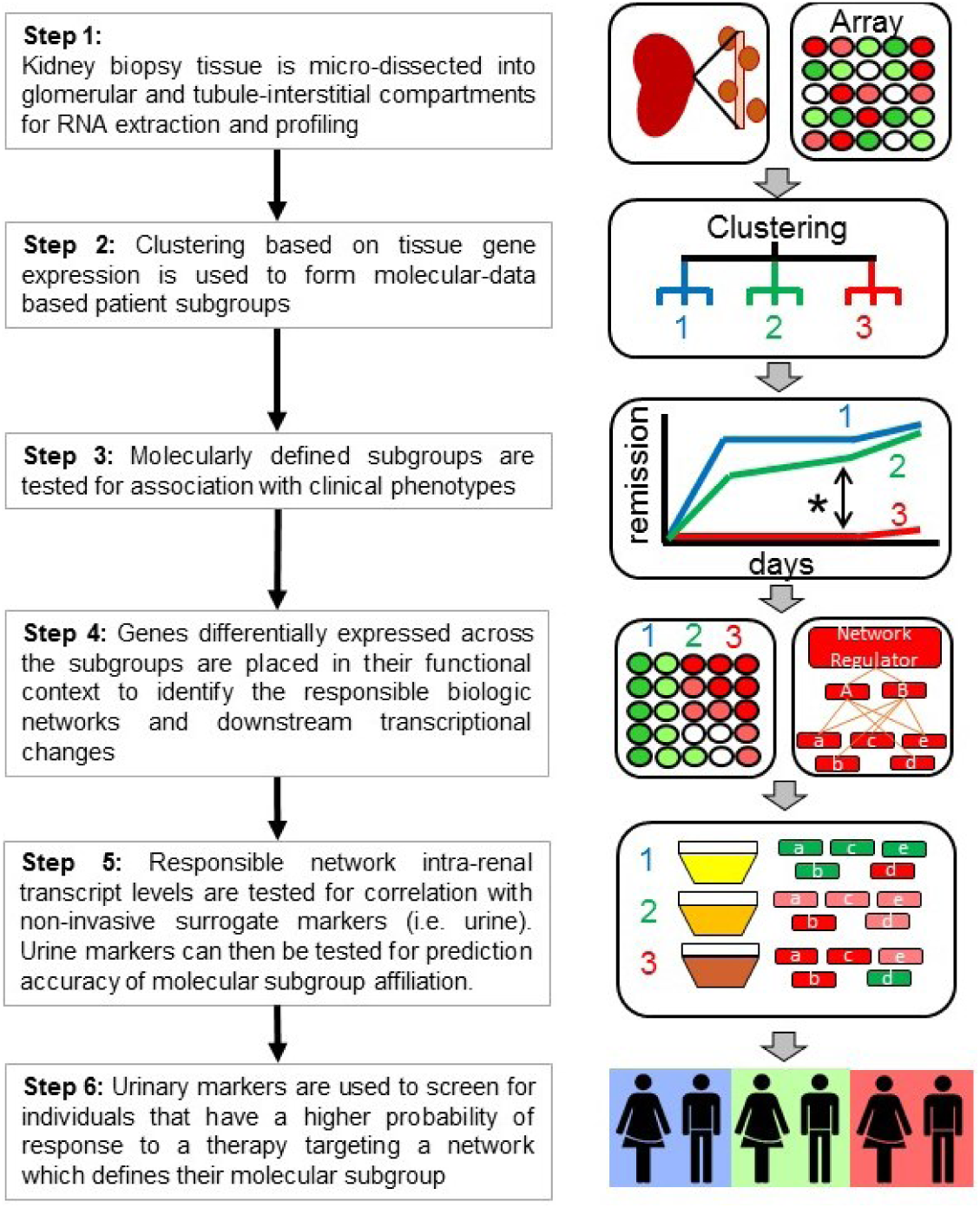
**Overall strategy** to identify non-invasive urinary markers for tissue-derived molecular patient subgroups.

Kidney diseases are uniquely positioned to implement this approach as a kidney biopsy is the diagnostic gold standard for NS, allowing for identification of molecular tissue signatures which can be linked to detailed histopathology assessment and non-invasive urine markers and validated against clinical outcome. In this study, we utilize the prospective Nephrotic Syndrome Study Network (NEPTUNE) cohort and its European sister network (European Renal cDNA Bank (ERCB)) to implement this approach and identify a sub group of patients with a shared molecular signature, potentially amenable to targeted therapy.

## Results

*Unbiased Hierarchical Clustering of Tubulointerstitial Compartment Gene Expression to Identify Molecular Subgroups of Nephrotic Syndrome:* 123 NEPTUNE patients with MCD and FSGS were clustered into three groups (n=62, 42, and 19, respectively) according to their tubulointerstitial mRNA expression levels from their clinically-indicated renal biopsy (Supplemental Figure 1). Baseline characteristics of the participants in each cluster are listed in table 1. Patients in cluster three were older, and had lower eGFR and higher UPCR at baseline. There was no difference in race, sex or duration of disease across the clusters. Although cluster 3 had a greater proportion with FSGS, all three clusters had participants with both MCD and FSGS according to the conventional histopathologic classification. In an unadjusted survival model, Cluster 3 had a more progressive phenotype, with lower hazard of complete remission (p-value 0.002) and greater hazard of the composite of ESRD or 40% decline in eGFR from baseline (p-value 0.007, Figure 2).

**Figure 2:**
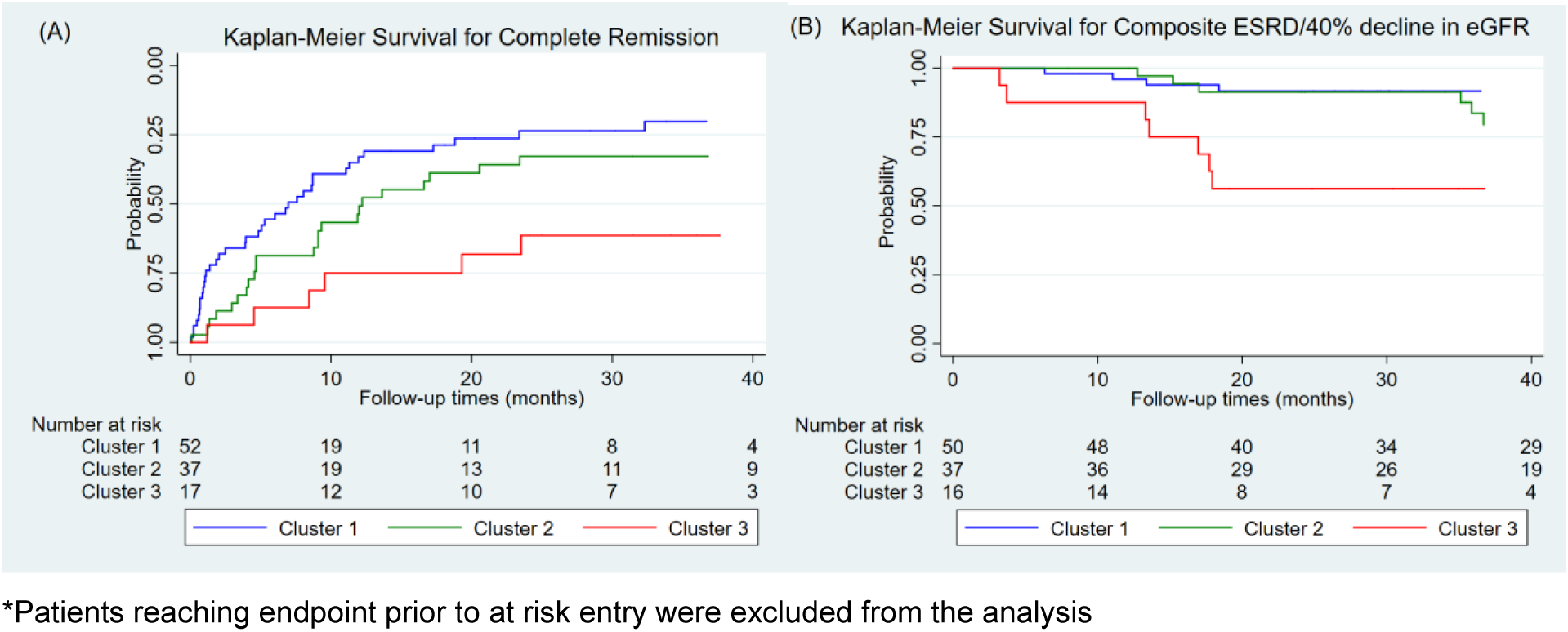
**Unadjusted Kaplan Meier curves by cluster membership** for complete remission of proteinuria from time of screening, p-value 0.002 (A), and composite of ESRD/40% drop in eGFR from baseline, p-value 0.007 (B).

**Table 1:** Baseline characteristics of participants by gene expression cluster. Continuous, normally distributed variables are presented as mean (SD). Continuous, non-normally distributed variables are presented as median (IQR). Categorical variables are presented as n(%).

|  | All<br>(N = 123) | Cluster 1<br>(N = 62) | Cluster 2<br>(N = 42) | Cluster 3<br>(N = 19) | p-value |
| --- | --- | --- | --- | --- | --- |
| **Age (years)** | 29 (22) | 17 (16) | 39 (23) | 44 (18) | <0.001 |
| Black Race | 40 (34%) | 18 (31%) | 12(30%) | 10 (53%) | 0.17 |
| Female | 41 (35%) | 22 (38%) | 18 (44%) | 6 (32%) | 0.31 |
| FSGS | 67 (57%) | 22 (38%) | 30 (73%) | 15 (79%) | <0.001 |
| Disease Duration<br>(months) | 4 (1, 26) | 4.5 (1.5, 17.5) | 6.5 (2, 52) | 1.0 (0, 20) | 0.06 |
| eGFR<br>(mL/min/1.73m2) | 88 (36) | 108 (31) | 79 (28) | 45 (20) | <0.001 |
| UPCR (mg/mg) | 1.2 (0.3, 3.5) | 0.7 (0.1, 2.7) | 1.5 (0.7, 3.6) | 2.4 (1.5, 4.6) | 0.01 |
| % IF | 5 (1, 19) | 1 (0, 5) | 9 (4, 19) | 27 (18, 56) | <0.001 |
| On RAAS Blockade | 67 (57%) | 24 (41%) | 31 (76%) | 12 (63%) | 0.003 |
| On IST | 60 (51%) | 37 (64%) | 18 (44%) | 5 (26%) | 0.01 |

\*eGFR: estimated glomerular filtration rate; MCD: Minimal Change Disease; FSGS: Focal Segmental Glomerulosclerosis; UPCR: Urine protein to creatinine ratio; IF: Interstitial Fibrosis; RAAS: Renin-angiotensin Aldosterone System; IST: Immunosuppressive Therapy

*Functional context of differentially expressed genes and replication in independent cohort:* 2517 genes were differentially regulated in the NEPTUNE cohort between cluster 3 versus 1 and 2, with a 1.5 fold-change and q-value <0.05. TNF itself was increased and found to center one of the top gene interaction networks from the differentially expressed gene set (Figure 3A). Genes were further analyzed to determine functional context of elevated TNF expression in Cluster 3. The canonical signal transduction pathways with the highest enrichment score was granulocyte adhesion and diapedesis with 54 of 151 (35.8%) pathway genes differentially expressed in cluster 3 (p-value<0.001). Differentially expressed genes in this pathway included TNF, which was one of the pathway activation inputs. In upstream regulator analysis (an analysis that takes into account both enrichment of and underlying direction of differential gene expression changes using cause and effect relationships), the top predicted activated protein network was TNF (IPA Z-score=10.2, enrichment p-value=3.65E-84, Figure 3B). A mechanistic network centered on downstream effects of TNF activation explained 26% (660/2517) of the differentially expressed genes in the analysis and included multiple transcription factors previously implicated in chronic kidney diseases including activation of the NFκB complex (as well as activation of NFKB1 (p105/p50) and RELA (p65) subunits) (*8–10*), and STAT1 and STAT3 (*11*). Lastly, 11 of the genes in the TNF causal network (including *TNF*) were supported by multiple literature assertions in IPA (Figure 3C), and were also profiled on a targeted proteomic profile panel.

**Figure 3.**
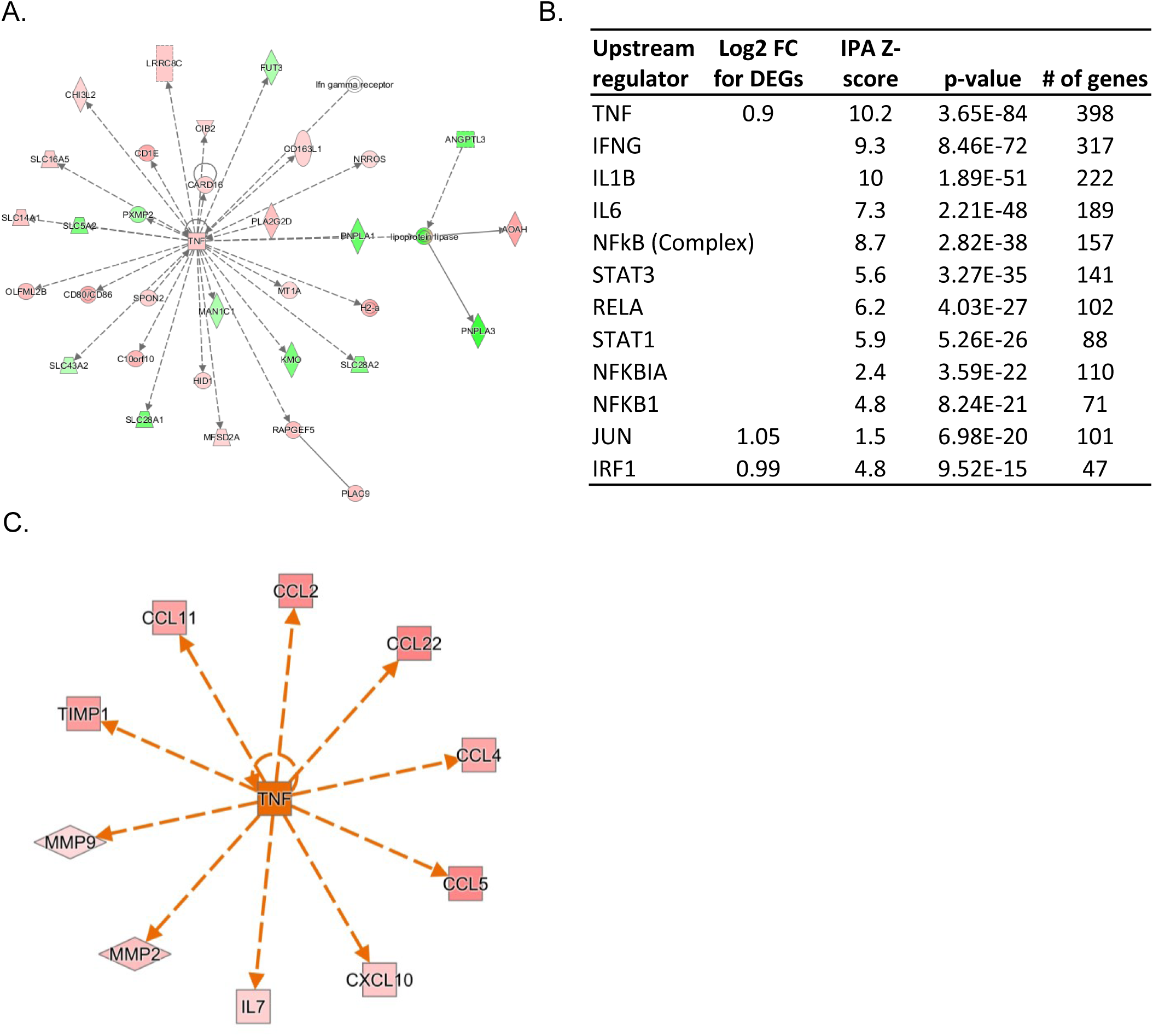
Transcriptomic profiles of subjects in cluster 3 compared to subjects in cluster 1 and 2 support TNF activation. **(A)** IPA gene interaction network centered on TNF. Genes up-regulated are colored in red and genes down-regulated are colored green **(B)** Upstream regulators that were part of the predicted TNF activation network. The number of genes supporting the IPA Z-score and p-value enrichment are shown. **(C)**. Genes causally downstream of the TNF activation network that were also part of the NEPTUNE biomarker panel. Orange indicates predicted activation of TNF and red genes indicate a gradient of fold change up-regulation in cluster 3 NEPTUNE participants.

To validate the molecular profiles identified in this cluster, unsupervised hierarchical clustering was applied to an independent cohort. Tubulointerstitial transcriptome data from 30 patients with MCD and FSGS in the European Renal cDNA Bank (ERCB) was used for validation (Supplementary Table 1). As in the NEPTUNE discovery cohort and three clusters were also identified. Patients in the ERCB cluster 3 also had significantly lower mean eGFR (35 ± 17, n=6, p<0.001) compared to the other two clusters (94±35 for the combined cluster 1 and 2, also see Supplementary Table 1). A differential expression analysis was also performed between cluster 3 and the other clusters. Genes that met a q-value<0.05 threshold in both NEPTUNE and ERCB displayed a high correlated expression profile (Supplemental Figure 2, R2 of Log2FC =0.84, p<0.001, with 93% of transcripts sharing directionality of change). To further validate the NEPTUNE findings, the same differential expression filter was applied (1.5 fold-change and q-value<0.05) to the ERCB cohort cluster 3 signature. This resulted in 703 genes and consistent with the findings from the NEPTUNE cohort, the top predicted protein network was activation of TNF (IPA Z-score=7.2, p-value=1.9E-22). Predicted activation of TNF explained 23% (163/703) of the differentially expressed genes in this cohort.

*Patient-level TNF score and relationship with cluster:* To be able to quantify TNF activation within individual patient samples, and assess its association with NS cluster assignment, a TNF activation score was generated using causal assertions were associated with predicted TNF activation in the NEPTUNE cohort. Starting with 398 genes that contributed to predicted TNF activation, genes were limited to those with multiple (≥3) lines of curated literature evidence (to limit spurious associations), and then further to those up-regulated by TNF (as a majority of genes (>95%) contributing to predicted TNF activation were up-regulated). This reduced the set of TNF-regulated genes to 145 (Supplementary Table 2). First, Log2 gene expression data for the 145 genes were converted to Z-scores across the NEPTUNE transcriptomic dataset. Next, the mean of each of the 145 Z-score gene expression values from each participant’s profile as the TNF activation score. Participants in cluster 3 had higher TNF activation scores than those in clusters 1 or 2. Mean (SD) score in cluster 3 was 1.01 (0.50), as compared to 0.01 (0.34) in cluster 2 and -0.53 (0.27) in cluster 1, p-value <0.01 (Figure 4). To address the potential for data overfitting, the 145 gene set was scored in a similar manner in the ERCB cohort. Consistent with differential gene expression profiles, and similar networks identified in the ERCB cohort, the association of TNF activation score with cluster 3 was confirmed in these samples (data not shown). Thus, a molecular signal consistent with TNF activation in primary NS was represented by a downstream gene signature in multiple cohorts.

**Figure 4:**
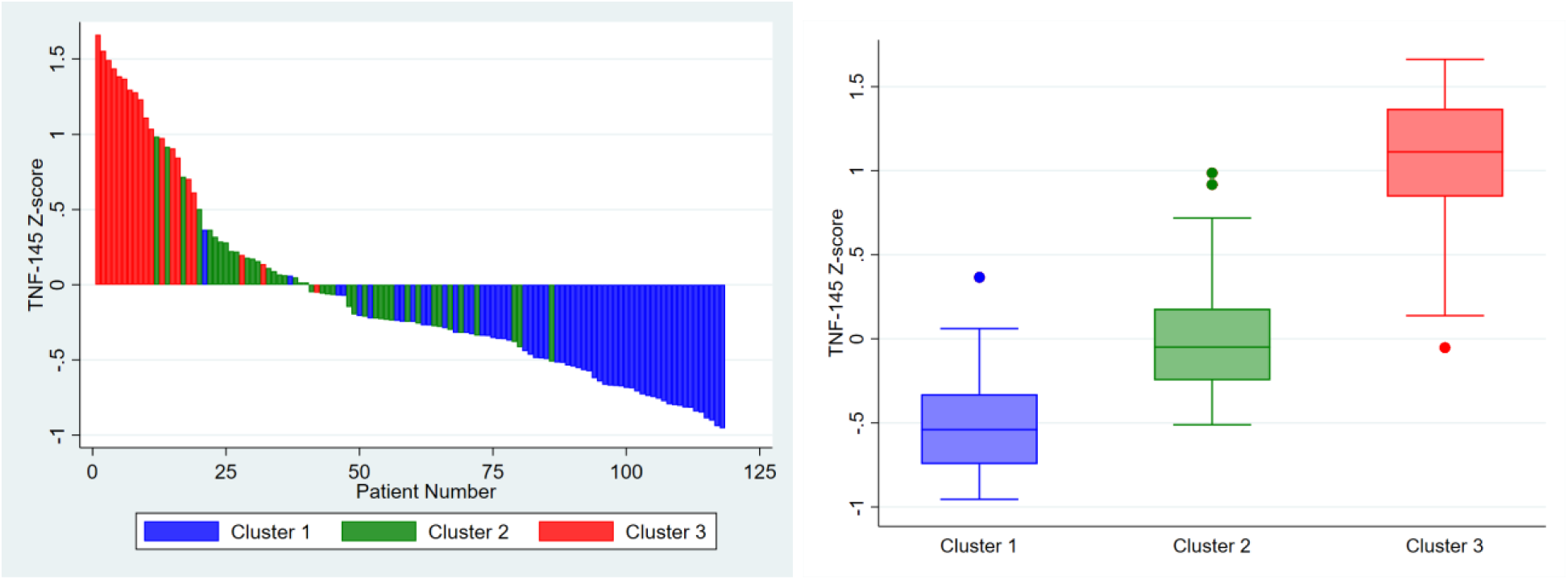
**Distribution of TNF patient scores** across all profiled participants and by cluster membership.

*Association of TNF activation score with clinical outcomes:* At baseline, TNF activation score was correlated with severity of interstitial fibrosis (rho = 0.69, p-value <0.001, Figure 5), but median (IQR) was 22.5 (10.5 - 49.5) and range was 0 to 71%. To evaluate to what extent TNF activation score from the renal tissue expression data captured the variability in loss of eGFR over time observed in cluster 3 as compared to clusters 1 and 2, a generalized estimating equation (GEE) model of eGFR over time was fit separately with cluster membership and TNF activation score as primary predictors of interest. After adjustment for demographics, diagnosis, time, baseline eGFR and UPCR, cluster 3 was associated with a 19 mL/min/1.73m2 lower eGFR during follow-up as compared to cluster 1. Cluster 2 was not significantly different from cluster 1. Similarly, in the fully adjusted model, a 1 unit greater TNF activation score was associated with a 12 mL/min/1.73 m2 lower eGFR during follow-up (Table 2).

**Figure 5:**
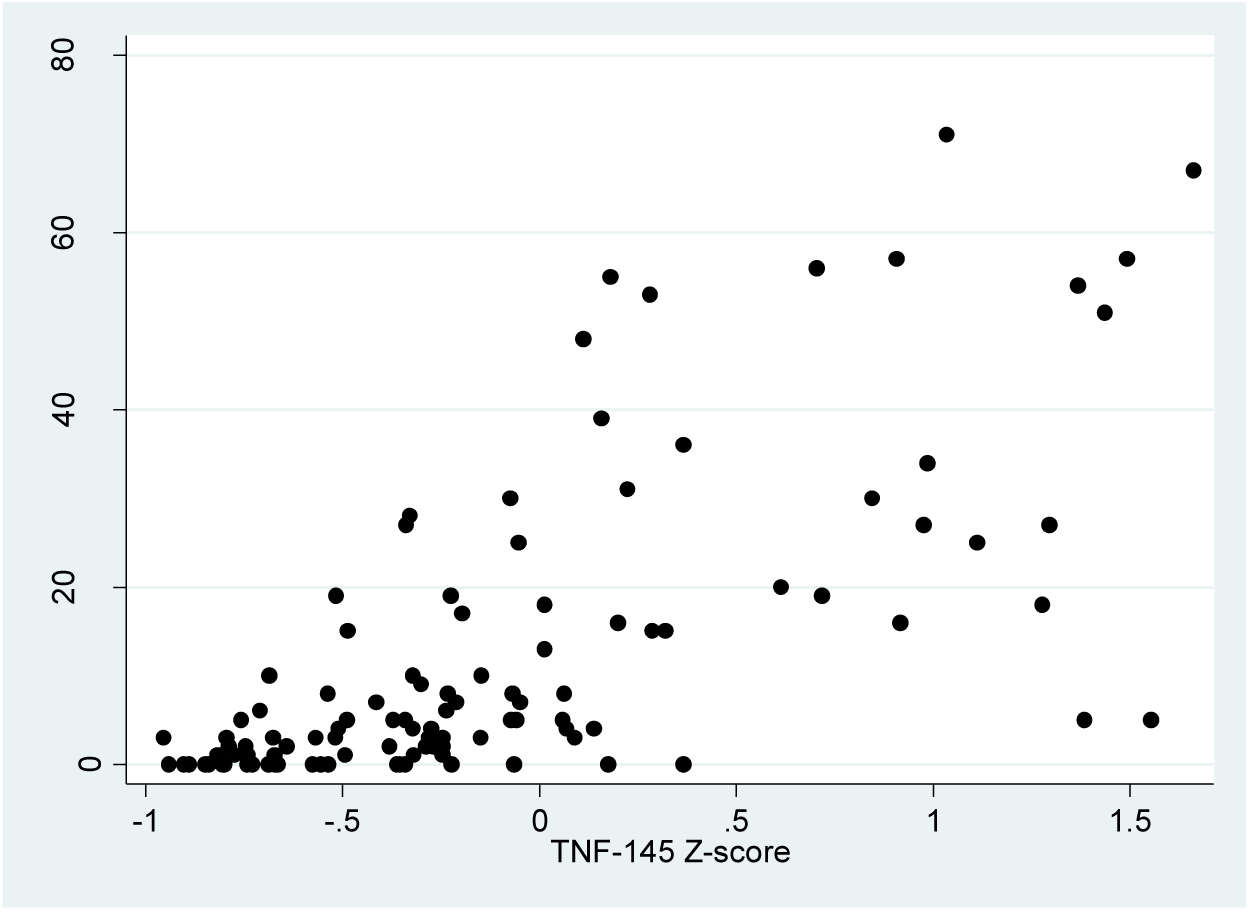
TNF alpha activation was correlated with interstitial fibrosis.

**Table 2:**
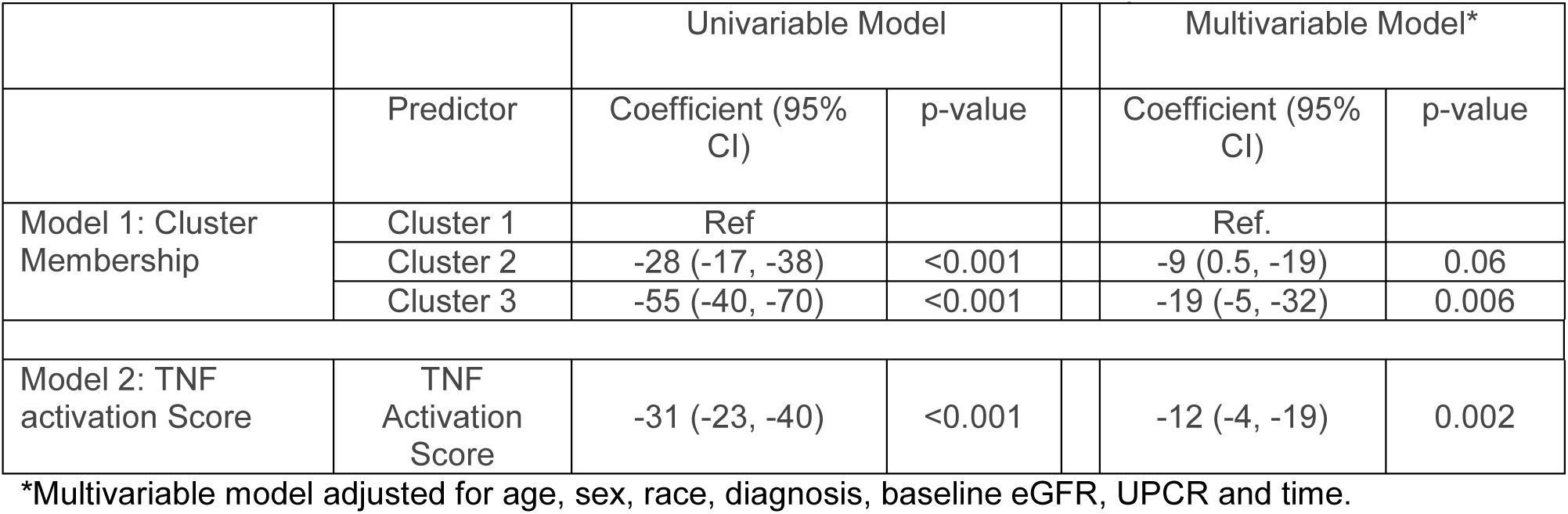
**Generalized Estimating Equations (GEE)** of eGFR (mL/min/1.73m2) after the baseline visit. Separate models for cluster membership and TNF activation score as primary predictors of interest.

*Non-invasive biomarker identification of TNF activation score:* Taking advantage of targeted proteomic data sets available as part of the multi-scalar data platform in NEPTUNE, profiles from 54 urinary cytokines, matrix metalloproteinases and tissue inhibitor of metalloproteinases were investigated. As shown in Figure 3C, 11 proteins with urine biomarker profiles were also part of a TNF causal network (i.e. gene expression values were downstream of TNF and a readout or signature of potential TNF activation in the kidney). Thus, we hypothesized that a biomarker or group of urine biomarkers might be sufficient to recapitulate intra-renal TNF activation and act as non-invasive surrogate biomarkers. Biomarkers with expression profiles in the dynamic range in at least 75% of samples, and those with a high level of intrarenal log2 mRNA versus log10 urine protein (normalized to creatinine) correlation (p<0.0001, r^2^≥0.25) in MCD and FSGS were chosen as potentially representative of the intra-renal transcriptional state (Figure 6A). Two genes, *CCL2* and *TIMP1* had corresponding urine proteomic profiles meeting these criteria (Figure 6B). Urine biomarker profiles for CCL2 (also known as MCP1) and TIMP1 were highly correlated with the TNF activation score (p<0.0001, r^2^≥0.25 for both biomarkers, Figure 6C). Thus, these biomarkers were identified as non-invasive surrogates reflective of the intra-renal transcriptional state and of the TNF activation score.

**Figure 6.**
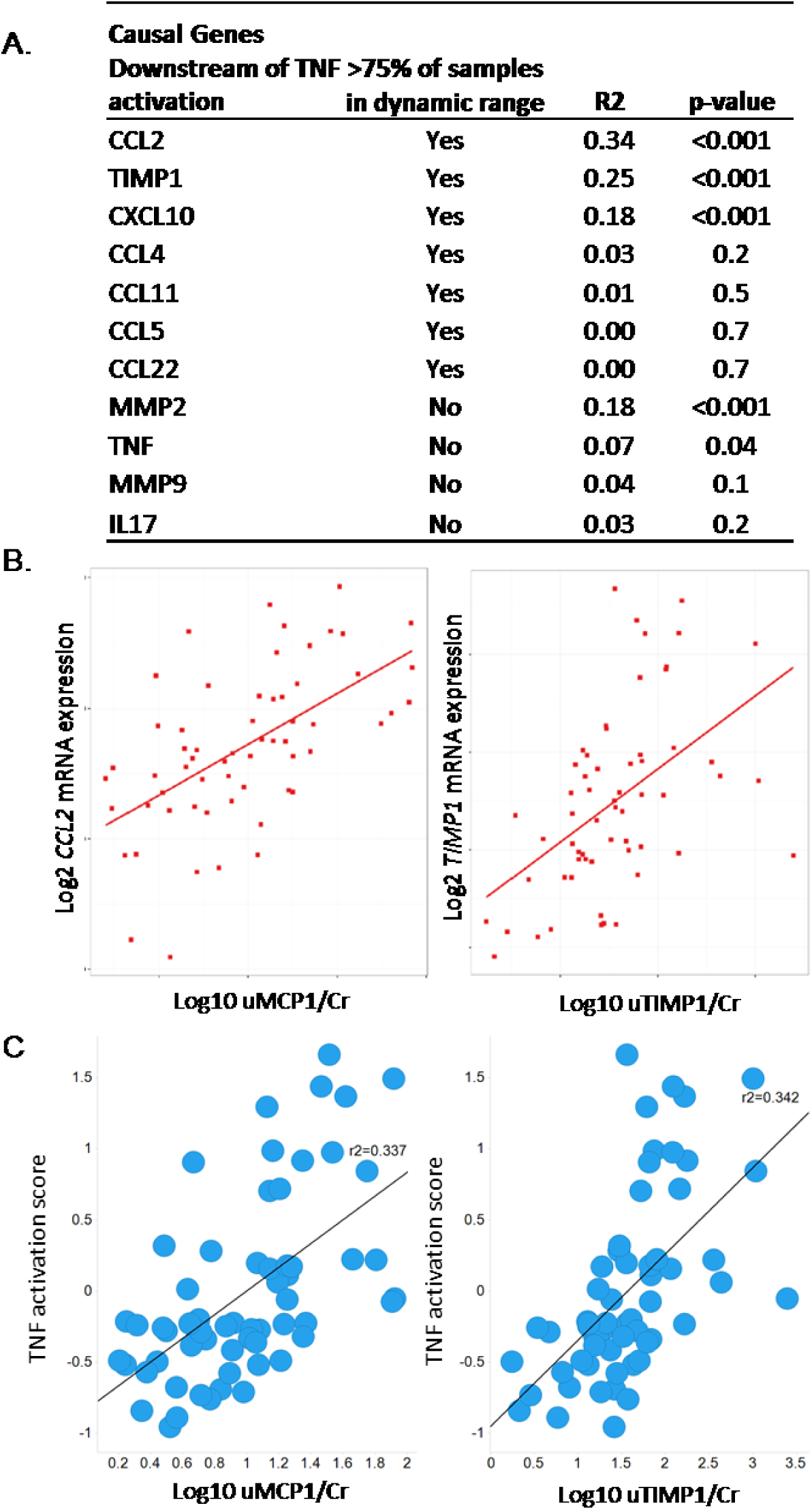
Urinary inflammatory biomarkers correlate with TNF activation score. **(A)** A prioritization schema was applied to identify biomarkers with the most reliable intra-renal mRNA and urine proteomic profile correlations. **(B)** Intra-renal and urine biomarker profile correlation plots in subjects with MCD or FSGS for CCL2 (left panel) and TIMP1 (right panel). **(C)** TNF activation score plotted against urine biomarker profiles for MCP1 and TIMP1.

*Predictive ability of biomarkers:* The base model presented here used eGFR and UPCR and additional models added the urinary biomarker levels of TIMP1 and MCP1. The fully adjusted model had highest c-statistic and positive predictive value for non-invasive assessment of the TNF activation score (Table 3).

**Table 3:**
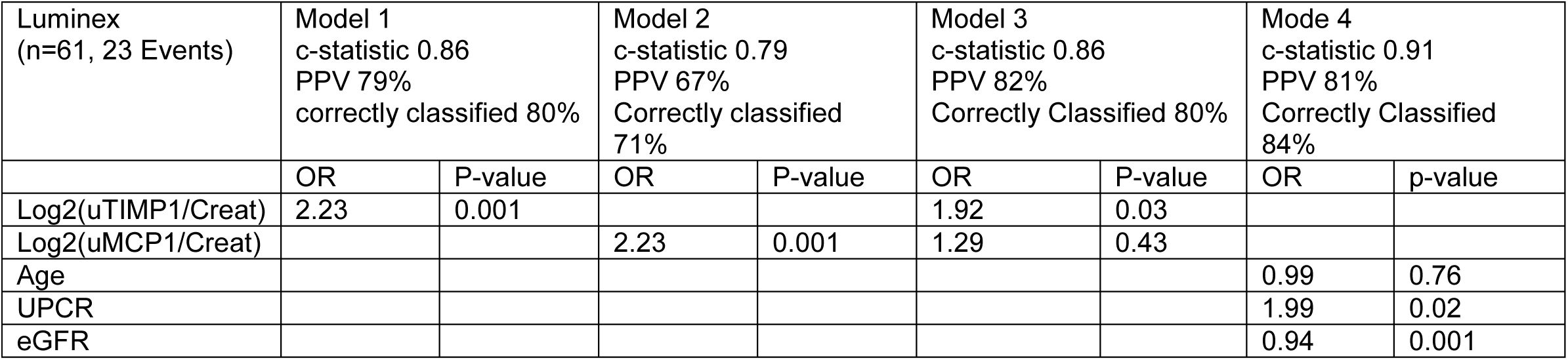
Logistic Regression of positive TNF activation score

## Discussion

This work introduces the concept of how a precision medicine strategy can work for nephrotic syndrome. The study utilized kidney biopsy tissue transcriptomics to identify a subgroup of nephrotic syndrome patients with a shared molecular profile and poor clinical outcome. Using an unbiased analysis of tubulointerstitial compartment gene expression data, without clinical or pathology data, a subgroup of participants was identified that had less remission of proteinuria and more loss of kidney function over time. The molecular profile of this group was evaluated for its underlying biological processes and found to center on TNF activation. TNF activation, quantified within individual patients, was sufficient to capture association with poor clinical outcome observed by cluster assignment. A combination of clinical features and urinary biomarkers could then be identified as non-invasive predictors of tissue TNF activation with high accuracy.

TNF is a pro-inflammatory, immunoregulatory cytokine, implicated in many systemic inflammatory diseases as well as kidney diseases (*12–14*). It is produced by infiltrating immune cells, but also by renal tissue cells, including podocytes and mesangial cells (*15*). In isolated rat glomeruli, TNF-alpha administration increased albumin permeability (*16*). In rats that spontaneously develop nephrotic syndrome and FSGS (Buffalo/Mna), renal expression of TNF increases before the onset of proteinuria (*17*). In humans, TNF levels from cultured peripheral blood mononuclear cells were higher in children with active nephrotic syndrome, compared to those in remission and controls (*18*). Case reports and small studies have reported that anti-TNF therapy may be effective in a subset of nephrotic syndrome patients, but no data was available on intra-renal activation of the pathway (*19–21*). Current clinical practice and diagnostic evaluation cannot identify this subset for targeted interventional trials.

Based on this animal and human evidence, the FONT trial (novel therapies in resistant FSGS) tested the TNF inhibitor adalimumab in patients with therapy-resistant FSGS using an unstratified approach (*4*). Of the total 16 patients treated in the phase I and phase II studies, 2 participants had dramatic improvements in proteinuria (from 17 to 0.6mg/mg and from 3.6 to 0.6 mg/mg in the other). Although the study is considered an unsuccessful trial in demonstrating efficacy of this therapy for all FSGS patients, a response in any of the patients with this severe phenotype is notable. This highlights the biologic heterogeneity underlying the recruited population to this study and similar clinical trials in FSGS. The observation of highly variable and unpredictable response to TNF blockade in the FONT trial is similar to that observed in routine clinical practice to standard therapies. It demonstrates the need for a precision approach to better assign patients to conventional therapies as well as offering access to experimental therapies in the setting of clinical trials to match patients to the most effective medication and sparing toxicity from unnecessary medications.

The pipeline described in this paper could be applied to clinical trial design whereby a nephrotic syndrome population could be enriched for patients with a higher probability of having a particular pathway upregulated and amenable to targeted therapy. Specifically, the coefficients from a validated logistic model could be used to calculate a probability of pathway activation as inclusion criteria for entry into a clinical trial. Thus, this would increase the chance that a trial would include a higher proportion of patients with a more homogeneous molecular profile amenable to the investigational target.

Several limitations of this approach are acknowledged. The clustering was done using the tubulointerstitial compartment as opposed to the glomerular compartment which is certainly also relevant to the pathophysiology of glomerular diseases. However, tubulointerstitial damage and fibrosis has been shown to be one of the strongest predictors of clinical outcome in the NEPTUNE cohort and treatment response (*22*), which crosses the conventional disease classifications. Medications targeting this common mechanism may be expected to have efficacy as disease modifying drugs in chronic kidney disease across multiple conventionally diagnosed renal diseases (*9*). For some patients, high TNF activation may represent a disease too advanced to be amenable to any therapy. However, the analysis did include samples from multiple patients with low interstitial fibrosis and high TNF scores. Bulk expression data was utilized and so differentially expressed genes may reflect differences in cell composition between the clusters. The accuracy of the non-invasive surrogates as dynamic, i.e. target engagement biomarkers requires validation and is being pursued in a proof of concept clinical trial under development.

In conclusion, this study implements a novel pipeline not previously applied in nephrotic syndrome patients to utilize tissue transcriptomics to identify a subgroup of patients with poor clinical outcomes. The potentially targetable pathway, TNF, was identified as a primary driver of disease and non-invasive markers could identify a patient population enriched for TNF activation. This mechanistic based disease classification is the first step to achieving the goal of assigning patients to therapies in a targeted manner and thus minimizing toxicity and maximizing benefit.

## Materials and Methods

*Study Participants:* The study was conducted on 123 participants with biopsy proven Minimal Change Disease (MCD) and Focal Segmental Glomerulosclerosis (FSGS) enrolled in the NEPTUNE study and who had tissue genome wide mRNA expression profiling completed. NEPTUNE is a multi-center, prospective study of children and adults with >500mg/day of proteinuria, recruited at the time of first clinically indicated baseline renal biopsy. Pathologic diagnosis is confirmed by review of digital whole slide images by study pathologists (*23*). Patients with evidence of other renal disease (e.g., lupus, diabetic nephropathy), prior solid organ transplant, and life expectancy < 6 months were excluded. The study enrolled at 21 clinical sites starting in August, 2010. The objectives and study design of NEPTUNE have been previously described (*24*) and can be found in the clinicaltrials.gov database under NCT1209000. Consent was obtained from individual patients at enrollment, and the study was approved by Institutional Review Boards of participating institutions. A subset of participants from the European Renal cDNA Cohort (ERCB) (n=30) with MCD and FSGS were used as a validation cohort for the gene expression analyses (*25*).

*Clinical Data:* NEPTUNE participants are followed prospectively, every 4 months for the first year, and then biannually thereafter for up to 5 years. Detailed information regarding socio-demographics, medical history and medication exposure are collected by subject interview and chart review. Local laboratory results are recorded and blood and urine specimens are collected at baseline and in each follow-up visit for central measurement of serum creatinine and urine protein/creatinine ratio. eGFR (mL/min/1.73m2) was calculated using the CKD-Epi formula for participants ≥18 years old and the modified CKiD-Schwartz formula for participants <18 years old. ESRD was defined as initiation of dialysis, receipt of kidney transplant or eGFR <15 mL/min/1.73m^3^ for two measurements. Complete remission was defined as UPCR <0.3 mg/mg on either a single void specimen or 24-hour urine collection. ERCB is a European multicenter study capturing renal biopsy tissue for gene expression profiling along with cross-sectional clinical information (e.g., demographics, eGFR) collected at the time of a clinically indicated renal biopsy(*25*).

*Molecular Data and Analysis*: Genome wide transcriptome analysis was performed on manually micro-dissected renal biopsy tissue that separated the tubulointerstitial compartment from the glomerular compartment. Total RNA was isolated, reverse transcribed, linearly amplified and hybridized on an Affymetrix 2.1 ST platform (NEPTUNE) and U133 platform (ERCB) as described previously (9, *26–29*). Gene expression was normalized, log-2 transformed and batch corrected with Entrez Gene ID annotations. Only genes expressed 1 standard deviation above the negative control were considered to be expressed and included in the analysis. Unsupervised hierarchical clustering and differential gene expression analysis was performed with Multiple Experiment Viewer (WebMeV, mev.tm4.org) using the tubulointerstitial compartment expression data. Differentially expressed genes between clusters of interest were analyzed for enrichment of canonical pathways and functional groups using the Ingenuity Pathway Analysis Software Suite (IPA).

*TNF Score:* Genes causally linked downstream of TNF were selected to compose a TNF activation score.(*30*) Selected genes were significantly up-regulated (≥1.5-fold change and q<0.05) in the differential expression gene set in cluster 3 compared to 1 and 2 and were predicted to be activated by TNF from 3 independent lines of evidence (i.e. literature references supporting the relationship) from IPA. 145 genes met these criteria (Supplemental table 3) and a z-score was generated for each gene for each patient. The individual z-scores across all 145 genes were averaged to calculate the composite TNF alpha activation score for each patient.

*Urine Biomarkers:* A panel of 54 urinary cytokines, matrix metalloproteinases and tissue inhibitor of metalloproteinases was available on a subset of NEPTUNE participants using the multiplex Luminex platform. All urine protein levels were normalized to urine creatinine. To be evaluated as a potential non-invasive marker of TNF activation, the urine protein had to be a product of a gene causally linked downstream of TNF and to be correlated with intra-renal tissue gene expression and TNF activation score.

*Statistical Analysis of the Association with Clinical data and Urine Biomarkers:* Descriptive statistics, including mean and standard deviation (SD) for normally distributed variables, median and interquartile range (IQR) for skewed variables and proportions for categorical variables were used to characterize baseline participant characteristics by molecular cluster. Multi-variable linear generalized estimating equations (GEE) were used to assess association of molecular cluster and TNF score with eGFR during follow-up. Pearson’s correlation was used to assess the relationship between TNF score and urinary biomarker concentration or mRNA expression. Urinary biomarker levels were divided by urinary creatinine to correct for urinary concentration/dilution and were log2 transformed to achieve a normal distribution. Logistic regression models were fit to assess the association of urinary biomarkers with a positive vs. negative TNF score. C-statistics were calculated from the logistic models to characterize the discrimination of the models. The improved predictive value of urinary biomarkers was assessed using the LR test for nested models. Analyses were performed using STATA, v12.1 (College Station, TX) with two-sided tests of hypotheses and p-value <0.05 as the criterion for statistical significance.

## Acknowledgement

The Nephrotic Syndrome Study Network Consortium (NEPTUNE), U54-DK-083912, is a part of the National Institutes of Health (NIH) Rare Disease Clinical Research Network (RDCRN), supported through collaboration between the Office of Rare Diseases Research, National Center for Advancing Translational Sciences and the National Institute of Diabetes, Digestive, and Kidney Diseases. Additional funding and/or programmatic support for this project has also been provided by the Else Kröner-Fresenius Foundation (ERCB), University of Michigan, the NephCure Kidney International and the Halpin Foundation, and the Applied Systems Biology Core at the University of Michigan George M. O’Brien Kidney Translational Core Center.

## Members of the Nephrotic Syndrome Study Network (NEPTUNE)

NEPTUNE Enrolling Centers

*Case Western Reserve University, Cleveland, OH:* J Sedor*, K Dell**, M Schachere^#^, J Negrey

*Children’s Hospital, Los Angeles, CA:* K Lemley*, L Whitted^#^

*Children’s Mercy Hospital, Kansas City, MO:* T Srivastava*, C Haney^#^

*Cohen Children’s Hospital, New Hyde Park, NY:* C Sethna*, K Grammatikopoulos^#^, R Odusayana

*Columbia University, New York, NY:* G Appel*, M Toledo^#^

*Emory University, Atlanta, GA:* L Greenbaum*, C Wang**, B Lee^#^

*Harbor-University of California Los Angeles Medical Center:* S Adler*, C Nast*‡, J La Page^#^

*John H. Stroger Jr. Hospital of Cook County, Chicago, IL:* A Athavale*

*Johns Hopkins Medicine, Baltimore, MD:* A Neu*, S Boynton^#^

*Mayo Clinic, Rochester, MN:* F Fervenza*, M Hogan**, J Lieske*, V Chernitskiy^#^

*Montefiore Medical Center, Bronx, NY:* F Kaskel*, N Kumar*, P Flynn^#^

*NIDDK Intramural, Bethesda MD:* J Kopp*, E Castro-Rubio^#^, J Blake^#^

*New York University Medical Center, New York, NY:* H Trachtman*, O Zhdanova**, F Modersitzki^#^, S Vento^#^

*Stanford University, Stanford, CA:* R Lafayette*, K Mehta^#^

*Temple University, Philadelphia, PA:* C Gadegbeku*, D Johnstone**, S Quinn-Boyle

*University Health Network Toronto:* D Cattran*, M Hladunewich**, H Reich**, P Ling^#^, M Romano^#^

*University of Miami, Miami, FL:* A Fornoni*, L Barisoni*, C Bidot^#^

*University of Michigan, Ann Arbor, MI:* M Kretzler*, D Gipson*, A Williams^#^, R Pitter^#^

*University of North Carolina, Chapel Hill, NC:* V Derebail*, K Gibson*, S Grubbs^#^, A Froment^#^

*University of Pennsylvania, Philadelphia, PA:* L Holzman*, K Meyers**, K Kallem^#^, A Swensen^#^

*University of Texas Southwestern, Dallas, TX:* K Sambandam*, E Brown**, M Cruz^#^

*University of Washington, Seattle, WA:* A Jefferson*, S Hingorani**, K Tuttle**§, L Curtin^#^, S Dismuke^#^, A Cooper^#§^

*Wake Forest University, Winston-Salem, NC:* B Freedman*, JJ Lin**, S Gray^#^

*Data Analysis and Coordinating Center:* M Kretzler, L Barisoni, C Gadegbeku, B Gillespie, D Gipson, L Holzman, L Mariani, M Sampson, P Song, J Troost, J Zee, E Herreshoff, S Li, C Lienczewski, T Mainieri, M Wladkowski, A Williams, D Zinsser

*National Institute of Diabetes and Digestive and Kidney Diseases (NIDDK) Program Office:* K Abbott, C Roy

*The National Center for Advancing Translational Sciences (NCATS) Program Office:* T Urv, PJ Brooks

* Principal Investigator; ** Co-investigator; # Study Coordinator

‡ Cedars-Sinai Medical Center, Los Angeles, CA

§ Providence Medical Research Center, Spokane, WA

## Supplementary Materials

**Supplemental Table 1:**
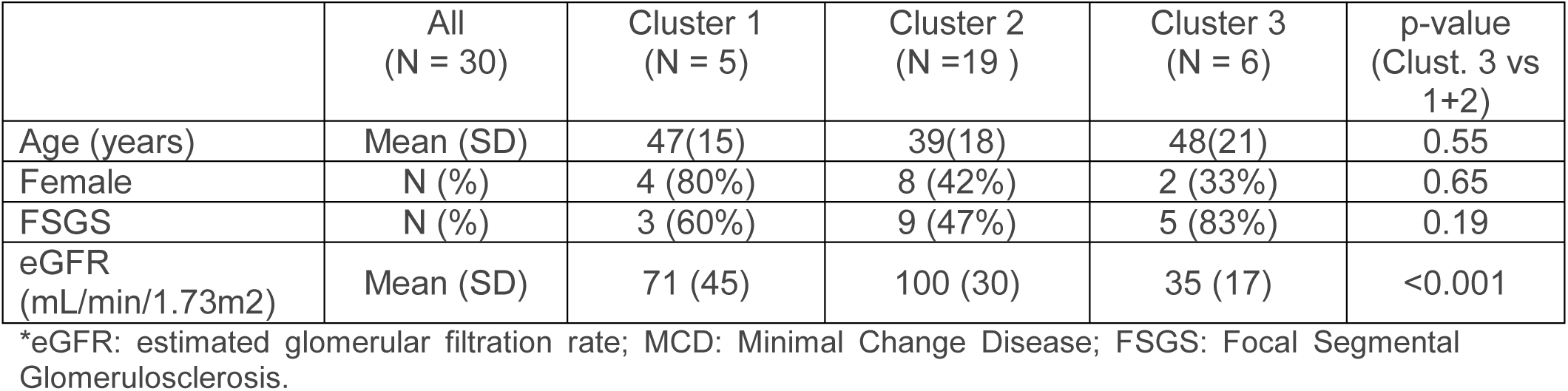
Baseline characteristics of ERCB participants by gene expression cluster. Continuous, normally distributed variables are presented as mean (SD). Continuous, non-normally distributed variables are presented as median (IQR). Categorical variables are presented as n(%).

|  | All<br>(N = 30) | Cluster 1<br>(N = 5) | Cluster 2<br>(N =19 ) | Cluster 3<br>(N = 6) | p-value<br>(Clust. 3 vs<br>1+2) |
| --- | --- | --- | --- | --- | --- |
| Age (years) | Mean (SD) | 47(15) | 39(18) | 48(21) | 0.55 |
| Female | N (%) | 4 (80%) | 8 (42%) | 2 (33%) | 0.65 |
| FSGS | N (%) | 3 (60%) | 9 (47%) | 5 (83%) | 0.19 |
| eGFR<br>(mL/min/1.73m<sup>2</sup>) | Mean (SD) | 71 (45) | 100 (30) | 35 (17) | <0.001 |

\*eGFR: estimated glomerular filtration rate; MCD: Minimal Change Disease; FSGS: Focal Segmental Glomerulosclerosis.

**Supplemental Table 2:** TNF-regulated genes contributing to the TNF activation score.

| Entrez ID | Gene Symbol | Entrez ID | Gene Symbol | Entrez ID | Gene Symbol | Entrez ID | Gene Symbol | Entrez ID | Gene Symbol |
| --- | --- | --- | --- | --- | --- | --- | --- | --- | --- |
| 12 | SERPINA3 | 1436 | CSF1R | 3575 | IL7R | 5359 | PLSCR1 | 7127 | TNFAIP2 |
| 31 | ACACA | 1520 | CTSS | 3587 | IL10RA | 5696 | PSMB8 | 7128 | TNFAIP3 |
| 154 | ADRB2 | 1524 | CX3CR1 | 3606 | IL18 | 5698 | PSMB9 | 7130 | TNFAIP6 |
| 165 | AEBP1 | 1536 | CYBB | 3624 | INHBA | 5699 | PSMB10 | 7133 | TNFRSF1B |
| 240 | ALOX5 | 1545 | CYP1B1 | 3627 | CXCL10 | 5788 | PTPRC | 7412 | VCAM1 |
| 241 | ALOX5AP | 1634 | DCN | 3659 | IRF1 | 5806 | PTX3 | 7424 | VEGFC |
| 330 | BIRC3 | 1848 | DUSP6 | 3676 | ITGA4 | 6036 | RNASE2 | 7474 | WNT5A |
| 355 | FAS | 1903 | S1PR3 | 3678 | ITGA5 | 6279 | S100A8 | 7852 | CXCR4 |
| 567 | B2M | 1906 | EDN1 | 3683 | ITGAL | 6288 | SAA1 | 7980 | TFPI2 |
| 597 | BCL2A1 | 1958 | EGR1 | 3684 | ITGAM | 6347 | CCL2 | 8870 | IER3 |
| 602 | BCL3 | 1999 | ELF3 | 3685 | ITGAV | 6351 | CCL4 | 9021 | SOCS3 |
| 629 | CFB | 2113 | ETS1 | 3689 | ITGB2 | 6352 | CCL5 | 9023 | CH25H |
| 718 | C3 | 2213 | FCGR2B | 3690 | ITGB3 | 6356 | CCL11 | 9180 | OSMR |
| 834 | CASP1 | 2335 | FN1 | 3694 | ITGB6 | 6363 | CCL19 | 9536 | PTGES |
| 837 | CASP4 | 2353 | FOS | 3725 | JUN | 6364 | CCL20 | 9636 | ISG15 |
| 920 | CD4 | 2634 | GBP2 | 3726 | JUNB | 6367 | CCL22 | 10512 | SEMA3C |
| 929 | CD14 | 2833 | CXCR3 | 3934 | LCN2 | 6372 | CXCL6 | 10537 | UBD |
| 942 | CD86 | 2920 | CXCL2 | 4050 | LTB | 6401 | SELE | 10563 | CXCL13 |
| 952 | CD38 | 3082 | HGF | 4071 | TM4SF1 | 6403 | SELP | 11221 | DUSP10 |
| 958 | CD40 | 3091 | HIF1A | 4233 | MET | 6422 | SFRP1 | 25816 | TNFAIP8 |
| 960 | CD44 | 3133 | HLA-E | 4313 | MMP2 | 6772 | STAT1 | 26298 | EHF |
| 1009 | CDH11 | 3383 | ICAM1 | 4318 | MMP9 | 6868 | ADAM17 | 51284 | TLR7 |
| 1026 | CDKN1A | 3428 | IFI16 | 4323 | MMP14 | 6890 | TAP1 | 58191 | CXCL16 |
| 1051 | CEBPB | 3459 | IFNGR1 | 4609 | MYC | 7040 | TGFB1 | 64332 | NFKBIZ |
| 1191 | CLU | 3489 | IGFBP6 | 4688 | NCF2 | 7052 | TGM2 | 79689 | STEAP4 |
| 1233 | CCR4 | 3554 | IL1R1 | 5054 | SERPINE1 | 7076 | TIMP1 | 112464 | PRKCDBP |
| 1236 | CCR7 | 3563 | IL3RA | 5284 | PIGR | 7097 | TLR2 | 114548 | NLRP3 |
| 1316 | KLF6 | 3566 | IL4R | 5328 | PLAU | 7099 | TLR4 | 414062 | CCL3L3 |
| 1435 | CSF1 | 3574 | IL7 | 5329 | PLAUR | 7124 | TNF | 729230 | CCR2 |

**Supplemental Figure 1:**
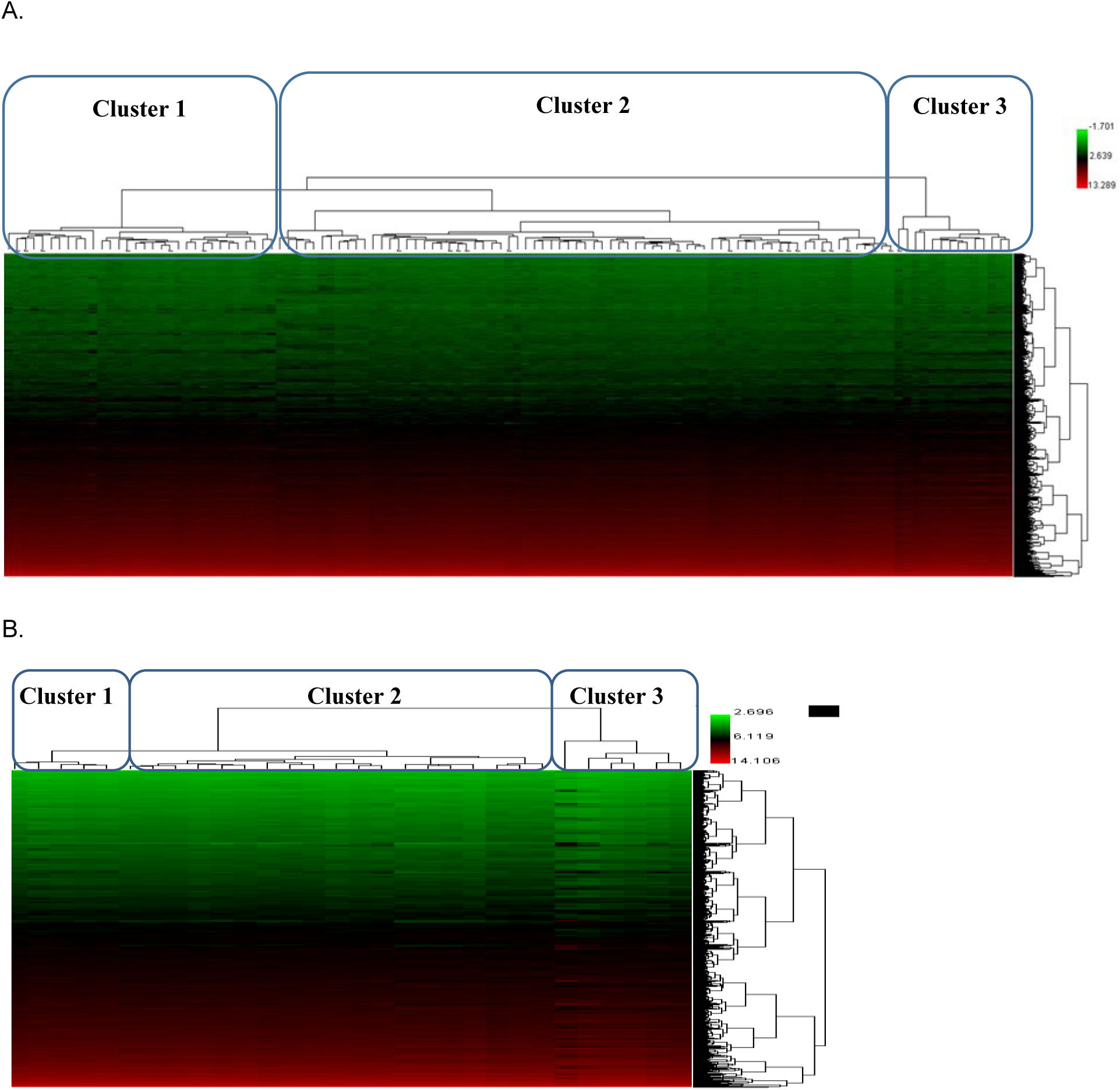
**Cluster dendrogram** of **(A)** NEPTUNE MCD and FSGS participants based on kidney biopsy tubulointerstitial gene expression data and **(B)** ERCB MCD and FSGS participants based on kidney biopsy tubulointerstitial gene expression data.

**Supplemental Figure 2.**
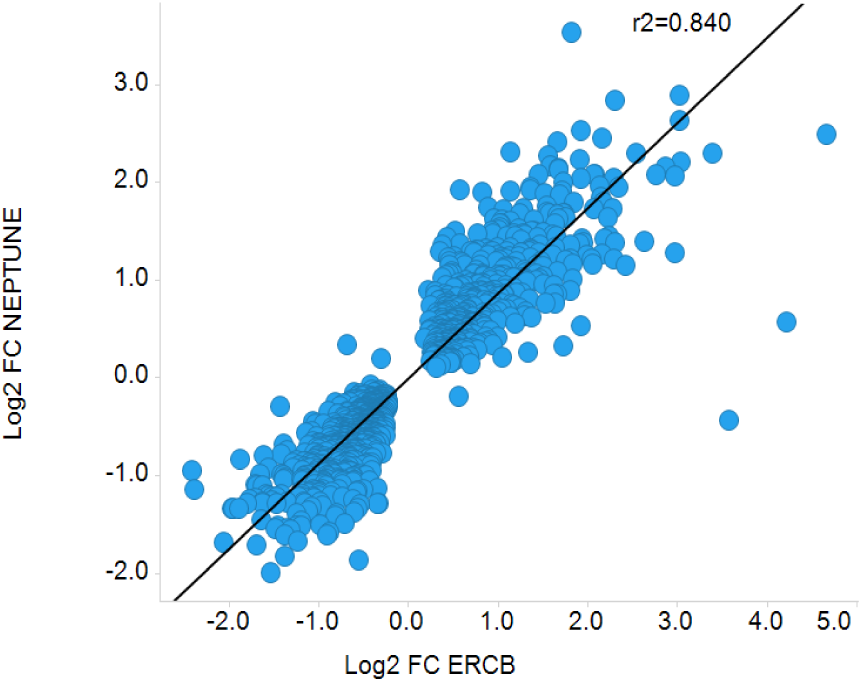
Log2 fold changes are presented for each dataset. Genes significantly differentially expressed (1,259 genes, q<0.05) in samples from cluster 3 patients for both NEPTUNE and ERCB cohorts are presented.

